# Understanding brewing trait inheritance in *de novo* lager yeast hybrids

**DOI:** 10.1101/2024.06.04.597363

**Authors:** Vasni Zavaleta, Laura Pérez-Través, Luis A. Saona, Carlos A. Villarroel, Amparo Querol, Francisco A. Cubillos

## Abstract

Hybridization between *Saccharomyces cerevisiae* and *Saccharomyces eubayanus* resulted in the emergence of *S. pastorianus*, a crucial yeast for lager fermentation. However, our understanding of hybridization success and hybrid vigour between these two species remains limited due to the scarcity of *S. eubayanus* parental strains. Here, we explore hybridization success and the impact of hybridization on fermentation performance and volatile compound profiles in newly formed lager hybrids. By selecting parental candidates spanning a diverse array of lineages from both species, we reveal that the Beer and PB-2 lineages exhibit high rates of hybridization success in *S. cerevisiae* and *S. eubayanus*, respectively. Polyploid hybrids were generated through rare mating techniques, revealing a prevalence of triploids and diploids over tetraploids. Despite the absence of heterosis in fermentative capacity, hybrids displayed phenotypic variability, notably influenced by maltotriose consumption. Interestingly, ploidy levels did not significantly correlate with fermentative capacity, although triploids exhibited greater phenotypic variability. The *S. cerevisiae* parental lineages primarily influenced volatile compound profiles, with significant differences in aroma production. Interestingly, hybrids emerging from the Beer *S. cerevisiae* parental lineages exhibited a volatile compound profile resembling the corresponding *S. eubayanus* parent. This pattern may result from the dominant inheritance of the *S. eubayanus* aroma profile, as suggested by the over-expression of genes related to alcohol metabolism and acetate synthesis in hybrids including the Beer *S. cerevisiae* lineage. Our findings suggest complex interactions between parental lineages and hybridization outcomes, highlighting the potential for creating yeasts with distinct brewing traits through hybridization strategies.

## INTRODUCTION

Hybridization, a phenomenon where genomes from two different species merge, is a significant evolutionary force across various kingdoms such as fungi (Gabaldón, 2020), Plantae (Glombik et al., 2020), and Animalia (Adavoudi & Pilot, 2022), facilitating rapid adaptive evolution. This process, more frequent among closely related sympatric species with incomplete prezygotic isolation, profoundly impacts the genetic diversity of a population (Gilman & Behm, 2011). Hybrid genotypes can exhibit traits that are not simply intermediate between their parents. In some cases, hybrids demonstrate increased fitness, a phenomenon known as hybrid vigor or heterosis (Birchler et al., 2006; Gabaldón, 2020), which can lead to replace parental species by hybrid swarms (Gilman & Behm, 2011). Notably, changes in the hybridization frequency between sympatric species have been correlated with anthropogenic environmental changes. An example of this is the fermentation environment, where high sugar concentration and temperature restrictions facilitated the hybridization of different *Saccharomyces* species, leading to the replacement of the original parental species (Hutzler et al., 2023; Sipiczki, 2019; Winans, 2022).

The *Saccharomyces* genus is an iconic model system to study hybridization (Alsammar & Delneri, 2020). Many *Saccharomyces* species have outcrossed, generating a range of hybrids (Stelkens & Bendixsen, 2022). Notably, most *Saccharomyces* hybrids have been isolated from industrial environments, such as those involved in the fermentation of wine and beer, suggesting that hybridization in yeast is an efficient mechanism to thrive in challenging fermentative conditions (Smukowski Heil et al., 2018). The best-known example is *Saccharomyces pastorianus*, the yeast responsible for lager-pilsner beer production by fermenting at low temperatures and the most produced alcoholic beverage globally (Gibson et al., 2017). *S. pastorianus* resulted from the successful interspecies hybridization between *S. cerevisiae* and *S. eubayanus* (Libkind et al., 2011), showcasing hybridization’s adaptative benefits. While *S. cerevisiae* is a well-studied species widely used in wine and beer fermentation (Legras et al., 2018), *S. eubayanus* was only recently discovered, with its characteristics remaining unknown until 2010 (Libkind et al., 2011). In this way, hybrids combine the *S. eubayanus* cold tolerance (due to its mitochondrial inheritance) and the superior fermentation kinetics and sugar consumption capacity from *S. cerevisiae* (Gibson et al., 2017). Throughout the history of lager yeast, different bottleneck events led breweries to retain only two pure-type strains: the ‘Frohberg’ and ‘Saaz’ lineages. This significantly contributed to a decline in the diversity of lager beer yeast (Gallone et al., 2019).

Industrial lager beers are known for their generally plain and homogeneous aroma profiles (Gonzalez Viejo et al., 2019). However, innovative craft beers with unique profiles are increasingly capturing consumers attention (Jaeger et al., 2020; Nieto-Villegas et al., 2024) Novel hybrids have played a significant role in boosting the production of desired fruity and floral volatile compounds (VCs), such as higher alcohols, ethyl esters, and fatty acid esters, in beer and other fermented beverages(Krogerus et al., 2016; Mertens et al., 2015; Pérez et al., 2022; Turgeon et al., 2021). Additionally, certain phenolic and spicy VCs, such as 4-vinyl guaiacol (4-VG), which are typically absent in commercial lager strains due to being considered off-flavors (POF-), are normally produced by wild yeasts such as *S. eubayanus* (POF+) (Gallone et al., 2016; Mertens et al., 2019; Urbina et al., 2020). The utilization of novel hybrid strains using these POF+ strains could expand the beer’s flavor profile, providing more complex fruity and spicy characteristics (Burini et al., 2021; Gonzalez Viejo et al., 2019; Hittinger et al., 2018).

Novel hybrid fermentation traits have been associated with the *de novo* lager’s subgenome composition. In this sense, a recent study using only three *de novo* lager yeast hybrids revealed that higher ploidy levels resulted in higher production of distinct volatile compounds and sugar consumption levels (Krogerus et al., 2016). These could likely result from genomic interactions and greater expression levels of flavor-active encoding genes, impacting the unique aroma profile in lager yeasts (Gibson et al., 2013; Krogerus et al., 2016). However, our current understanding of the parental subgenomes’ contribution, dosage, and ploidy levels concerning fermentative capacity and volatile compound production in *de novo* hybrids during lager fermentation is still limited. In addition, as the exact *S. cerevisiae* and *S. eubayanus* parental genomes of *S. pastorianus* are unavailable, our understanding of the evolutionary history of the lager hybrid is based on the sequence analysis of reference genomes from the parental species (Nespolo, et al., 2020; Peter et al., 2018), or other lager hybrid genomes (Gallone et al., 2019; Langdon et al., 2019). Consequently, the complex molecular interactions and the mechanisms by which genome interactions impact the fermentation and aroma profiles of *S. cerevisiae* x *S. eubayanus* hybrids remain largely unknown.

In this study, we aimed to deepen our understanding of *S. cerevisiae* and *S. eubayanus* hybridization, specifically focusing on creating and evaluating a genetically rich set of hybrids for beer production. Our research explores into these hybrids’ fermentative and aroma characteristics, evaluating their efficiency compared to parental strains. Through transcriptome analyses of *de novo* lager hybrids, our goal was to unravel how specific genetic combinations influence these aroma profiles. This research contributes to the scientific knowledge of hybridization and explores new avenues for enhancing yeast strains in the biotechnology industry.

## METHODOLOGY

### Yeast strains

Ten diploid *S. eubayanus* strains representative of different genetic lineages isolated from native Chilean forests were chosen (Nespolo et al., 2020) (**Table S1**). All these strains previously exhibited the highest fermentative capacities within each clade (Nespolo et al., 2020). In addition, twenty diploid *S. cerevisiae* strains isolated from different anthropogenic niches were included in this study (Peter et al., 2018). The commercial lager strain *Saccharomyces pastorianus* W34/70 was used as a fermentation control.

### Wort Fermentations

Depending on the specific experiment, fermentations were conducted in 10 mL and 50 mL volumes, using 12 °Plato (°P) beer wort. The wort was oxygenated to 15 mg l^-1^ and supplemented with 0.3 ppm Zn^2+^ (as ZnCl_2_) at 12 °C as previously described (Molinet et al., 2022). Briefly, a pre-inoculum was prepared overnight in 5 ml of 6 °P malt extract (Maltexco, Chile) wort at 20°C, which was then used to inoculate 50 ml culture in 12 °P malt extract under the same previous conditions. We inoculated 50 ml of 12 °P malt extract with 9x10^8^ cells for fermentations. The micro-fermentations were conducted for 14 days at 12 °C, and the CO_2_ production was recorded daily by weighing the bottles. Residual sugars and ethanol production were measured using HPLC. For this purpose, 20 µl of each sample filtered through 0.22 µm syringe filters was injected into Shimadzu Prominence HPLC equipment (Shimadzu, USA) and eluted on an Aminex HPX87H column (Bio-Rad, USA) using H_2_SO_4_ 5 mM as mobile phase and acetonitrile 4 ml/l at a flow rate of 0.5 ml/min.

### Generation of polyploid hybrids by rare mating

Polyploid hybrids were generated using the rare mating procedure as previously described (Pérez et al., 2022). We selected natural auxotroph variants of *S. cerevisiae* and *S. eubayanus* by plating overnight cultures on two minimal plates containing 2% glucose, 0.17% YNB without amino acids and (NH_4_)_2_SO_4_ (Difco, France). *S. eubayanus* tryptophan auxotrophic variants were selected on minimal media supplemented with 0.05% w/v 5-fluoroanthranilic acid (5-FAA) (Sigma-Aldrich, USA) and an amino acid stock previously reported (Toyn et al., 2000). *S. cerevisiae* lysine auxotrophs variants were obtained on minimal media plates supplemented with 0.1% w/v α-aminoadipic acid (α-AA) (Alfa Aesar, USA) and 30 mg/L lysine (Zaret & Sherman, 1985). Separate overnight cultures in YPD at 25°C were combined, centrifuged, and incubated for 7 days at 12 °C in fresh YPD. Potential hybrids were plated on a defined minimal medium consisting of 0.17% YNB without amino acids (Difco, France), 2% glucose, and 2% agar and incubated at 12°C for 7-10 days. Hybrid strains were confirmed by *Hae*III (NEB, USA) digestion of the ITS amplicon obtained using ITS1 and ITS4 primers (Krogerus et al., 2016; White et al., 1990).

As we plated combined cultures multiple times to procure hybrid colonies, we determined the success rate by calculating the frequency of successful attempts. Each attempt encompassed plating the combined culture on batches comprising 10 minimal media plates. A positive attempt was defined as identifying at least one confirmed hybrid within a specific batch of plates.

### Hybrid’s genetic stabilization and ploidy level determination

The hybrid strains’ genetic stabilization was performed as described by Lairón-Peris et al. (2020) and Pérez et al. (2022), with minor modifications. Briefly, 20 mL of 12 °P beer wort was inoculated with each hybrid strain for 7 days at 12 °C. Following this, 50 µl was used to inoculate fresh medium under the same conditions. This cell transfer process was repeated seven times, corresponding to approximately 42 generations. After the stabilization, the ploidy level of each hybrid was determined using flow cytometry as described by Nespolo et al. (2020). A 5 ml overnight culture of each hybrid in YPD medium was prepared, then pelleted, and resuspended in 2 ml of water. Subsequently, 1 ml of the suspension was mixed with 2.3 ml of cold absolute ethanol and stored at 4 °C for 24 h for fixation. The cells were then pelleted and resuspended in 1 ml of 50 mM sodium citrate buffer (pH 7). This step was repeated, and 1x10^7^ cells contained in 100 µl of 50 mM sodium citrate buffer were treated with 1 µl of 100 mg/ml RNAse A (Roche, Suiza) and incubated for 2 h at 37 °C to remove RNA. Finally, 350 µl of the labeling solution containing 50 µg/ml propidium iodide (Sigma-Aldrich, USA), 50 mM sodium citrate, pH7, was added, followed by a 40-minute incubation in darkness at room temperature. Samples were analyzed using a FACSCanto II cytometer (Becton Dickinson, USA) and around 150.000 single labelled cells were used for ploidy determination.

### Microculture and temperature tolerance phenotyping

Cells were pre-cultivated at 20°C without agitation for 48 h in 96-well plates containing 200-μL YPD (1% yeast extract, 2% peptone, and 2% glucose). A volume of 10 μl of pre-inoculum was used to inoculate a new 96-well plate containing 200 μl of YNB 0.67% (Difco, France) supplemented with the following carbon sources: 2% maltotriose (Sigma-Aldrich), 2% maltose (SRL, India), 20% maltose, 2% maltose with 6, 8 or 12 %v/v ethanol to an optical density (OD600) of 0.03–0.1. Additionally, the strains were assessed in complex media such as YPD, 12 and 20 °P. wort. OD600 for each well was measured at 620 nm every 30 min for 96 h. The average Area Under the growth Curve (AUC) was calculated using the R based tool Growthcurver v 0.3.1 (Sprouffske & Wagner, 2016). Values were normalized between 0 and 1, representing the lowest and highest growth values under a specific condition.

A serial dilution assay was carried out to assess yeast growth across a broad temperature range. For this, overnight YPD cultures of each strain were serially diluted, and 4 μl of each dilution was transferred to YPD plates. Inoculated plates were incubated at 4, 12, 20, 25, 30, and 37 °C. Plates incubated at 25, 30, and 37 °C were photographed on the 3rd day. Plates incubated at 20, 12, and 4 °C were photographed on the 4th, 8th, and 12th day, respectively.

### Volatile compounds quantification

Volatile compounds in both the hybrids and parental strains were analyzed using headspace solid phase microextraction (HS-SPME) with a 100-μm polydimethylsiloxane (PDMS) fiber (Supelco, Sigma-Aldrich, Spain). The analysis was conducted using a TRACE GC Ultra gas chromatograph equipped with a flame ionization detector (FID) and a TriPlus RSH autosampler (Thermo Fisher Scientific, Waltham, MA). Samples derived from 14-day beer wort fermentations at 12 °C were prepared by mixing 5 ml of the sample with an equal volume of NaCl saline solution (75 g of NaCl in 247.5 ml mQ-water) in a 20 ml glass vial. Each sample was incubated at 40 °C for 30 min before injection. The PDMS fiber was exposed to the vial’s headspace for 15 min, followed by a 5-minute desorption at 250 °C in spitless mode in the GC-injection port. Helium was used as the carrier gas at a flow rate of 1 ml/min. An Agilent HP INNOWax capillary column (30 m × 0.25 m), coated with a 0.25-µm layer of cross-linked polyethylene glycol (Agilent Technologies, USA), was employed.

The oven temperature program began at 50 °C for 5 minutes, increased by 1.5 °C/min to 100°C, then by 3 °C/min to 215 °C, and held for 2 minutes at 215 °C. The detector temperature remained constant at 280 °C. Chromatographic signals were recorded using ChromQuest software. Compounds were identified and quantified based on retention times and calibration curves of corresponding standard volatile compounds.

### Estimation of Best Parent heterosis

To evaluate best parent heterosis (BPH) we calculated BPH values as previously described by Geng et al., (2021). For this, we used the formula BPH = (F1 - BP)/BP, where F1 represents the value of a specific phenotype in the hybrid, and BP corresponds to the value of the same phenotype in the best parent. For heatmap visualization, the BPH value for a specific trait was normalized relative to the highest absolute value recorded among the hybrids for that trait, establishing a scale from 0 to 1. Values in red (>0 to 1) represented heterosis, whereas zero and negative values (0 up to -1) shown in shades of blue and white, respectively, represented the absence of heterosis.

### RNA-Seq

RNA from hybrids was extracted using E.Z.N.A. Total RNA Kit 1 (Omega Bio-Tek, USA) and subsequently purified using the RNeasy MinElute Cleanup Kit (Qiagen, Germany). The cDNA libraries were generated using the TruSeq RNA Sample Prep Kit v2 (Illumina, San Diego, CA, USA). Illumina libraries from hybrids were generated using paired-end 150 bp reads on an Illumina Next seq 500 as previously described (Venegas et al., 2023). RNA sequencing data from HB6 and HB41 hybrid strains were mapped against a concatenated *S. cerevisiae* R64-1-1 and *S. eubayanus* genome CL216.1 (Mardones et al., 2020) using STAR (--outSAMmultNmax 1), after which gene counts were obtained using featureCounts (Dobin et al., 2013; Liao et al., 2014). Expression count data were imported into R, with gene identifiers updated by parental species-specific mappings (*S. cerevisiae* and *S. eubayanus*). Differential expression was assessed using the DESeq2 package, version 4.1.2 (Love et al., 2014), applying a log2 fold change threshold (log2FC) of |1| and an adjusted *p*-value threshold (p-adj) of <0.01. Differentially expressed genes (DEGs) were thus identified in all conditions, and DEGs from HB6 (p-adj <0.01 and log2FC > 1) and HB41 (p-adj <0.01 and log2FC < -1) were subsequently extracted for further analysis.

Metabolic pathway enrichment was analyzed using the enrichKEGG and enrichGO tools (Wu et al., 2021), based on gene identifiers in Entrez format. Analyses were performed separately for each hybrid strain. The results of KEGG pathway enrichment and GO terms were visualized using dot and bar graphs.

### Statistical analysis

The data were deemed statistically significant with a *p*-value < 0.05, calculated using One-way ANOVA (analysis of variance) and the nonparametric Wilcoxon–Mann–Whitney test. These analyses were conducted in R (version 4.3.1) using the aov and wilcox.test functions, respectively. Plots and heatmaps were also generated with the ggplot2 (version 3.4.3) and pheatmap (version 1.0.12) packages.

## RESULTS

### High hybridization rate across most S. cerevisiae and S. eubayanus lineages

To maximise the genetic diversity of *S. cerevisiae* x *S. eubayanus* hybrids, we employed a crossbreeding strategy using yeast strains derived from distinct genetic lineages of both species. Initially, we aimed to identify parental strains with robust fermentative performance in beer wort at low temperatures. We evaluated 30 diploid strains, including 10 *S. eubayanus* and 20 *S. cerevisiae* strains, representing various lineages. The selection of *S. eubayanus* strains was based on their previously reported high fermentative capacity (Nespolo, et al., 2020). In agreement with our previous findings, no significant variations in fermentative capacity across *S. eubayanus* were observed (*p*-value > 0.05, one-way ANOVA, **Figure 1A**), and none of them exhibited maltotriose consumption (**Table S2A**). Notably, the fermentative capacity of these strains closely resembled that of the commercial strain W34/70 (*p*-value > 0.05, one-way ANOVA, **Figure 1A**). In contrast,

**Figure 1.**
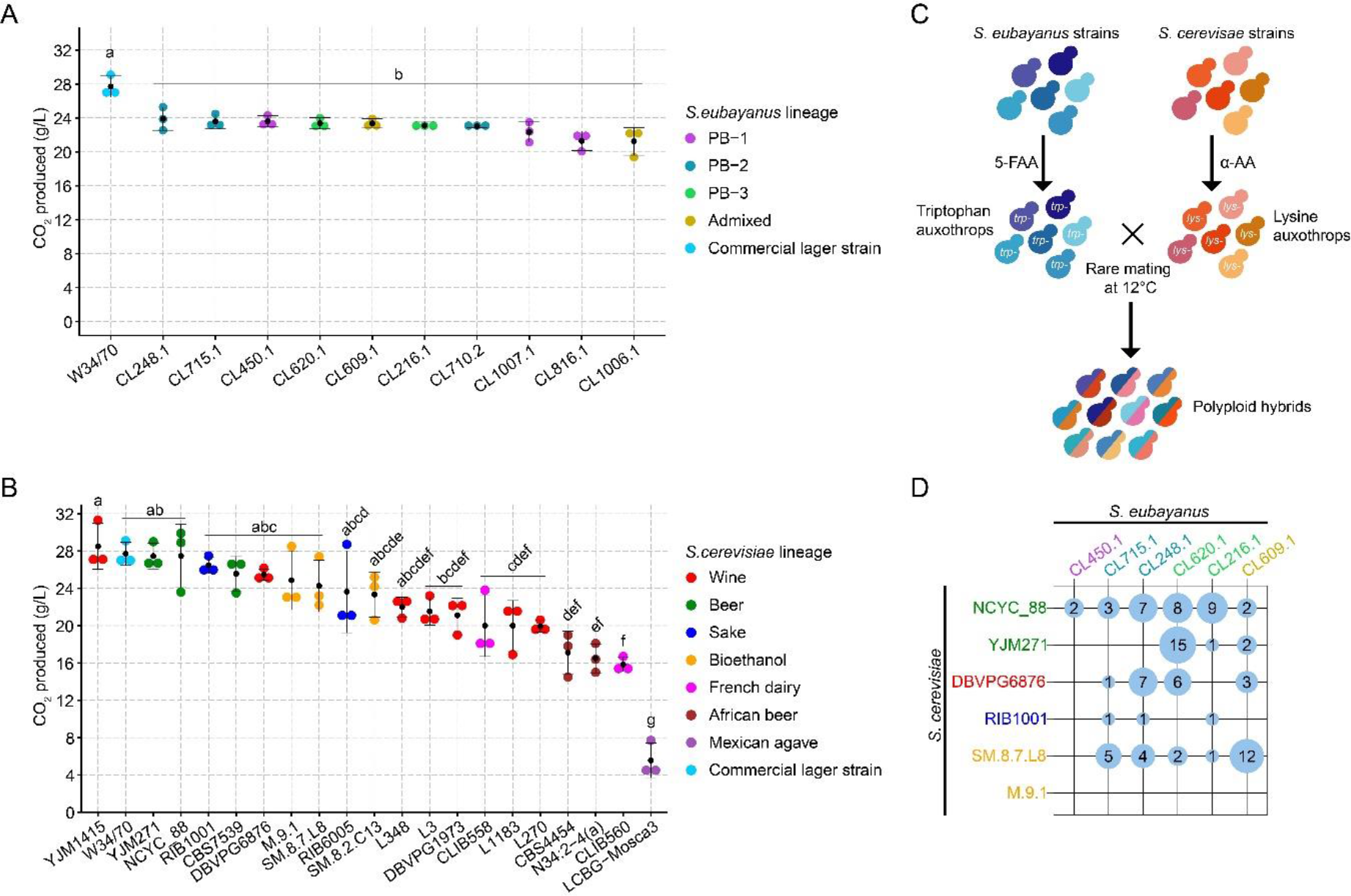
Hybridization between *S. cerevisiae* and *S. eubayanus* strains. (A) Total CO_2_ production (g/L) in (A) *S. eubayanus* and (B) *S. cerevisiae* strains under 12 °P wort from a broad range of lineages in both species. Black dots depict mean values between the three replicates. (C) Hybridization strategy to generate *de novo* polyploid lager hybrids using a rare mating approach. Tryptophane (*tyr-)* and lysine (*lys-*) auxotrophs were generated in *S. eubayanus* and *S. cerevisiae* under 5-fluoroanthranilic acid (5-FAA) and α-aminoadipic acid (α-AA), respectively. Subsequently, rare mating at 12°C was performed, and hybrids were selected using minimal media. (D) Hybridization success rate between *S. cerevisiae* and *S. eubayanus* strains. Different letters reflect statistically significant differences between strains with a *p*-value < 0.05, one-way ANOVA.

*S. cerevisiae* strains showed significant differences in fermentative capacity (*p*-value < 0.05, one-way ANOVA, **Figure 1B**). Strains from the Beer, Sake, Bioethanol, and specific strains from the Wine lineages displayed the highest fermentative capacity, with no significant differences compared to W34/70 (**Figure 1B**). Remarkably, six strains—YJM271 (Beer), NCYC_88 (Beer), YJM1415 (Wine), CBS7539 (Beer), RIB1001 (Sake), and DBVPG6876 (Wine)—exhibited maltotriose consumption exceeding 60% (**Table S2B**). Strains from the African Beer, Mexican Agave, French Dairy, and most of the Wine/European lineages showed a lower fermentative capacity than the commercial strain (*p*-value < 0.05, one-way ANOVA, **Figure 1B**). Based on the fermentative capacity and sugar consumption profiles, we selected six representative *S. eubayanus* strains from different lineages (CL248.1 (PB-2), CL715.1 (PB-2), CL450.1 (PB-1), CL620 (PB-3), CL609.1

(Admixed), and CL216.1 (PB-3)) and seven *S. cerevisiae* strains (YJM1415 (Wine), YJM271 (Beer), NCYC_88 (Beer), RIB1001 (Sake), DBVPG6876 (Wine), M9.1 (Bioethanol), and SM.8.7.L8 (Bioethanol)), all characterized by higher fermentative capacity from each parental species, and maltotriose consumption levels in *S. cerevisiae*. These chosen strains represented the genetic starting material to generate *S. cerevisiae* x *S. eubayanus* hybrids.

To generate polyploid hybrids, we employed a rare mating strategy involving crossbreeding complementary amino acid auxotrophic individuals from both species (**Figure 1C**). We successfully obtained auxotrophic colonies for both species, except for the *S. cerevisiae* YJM1415 wine strain. Each auxotrophic strain was subjected to crossbreeding at low temperatures (to facilitate the selection of hybrids containing the *S. eubayanus* mitochondria) with its counterpart species in liquid media. Among the 36 potential cross combinations, 21 produced at least one positive hybrid colony (**Figure 1D**). A total of 93 hybrid colonies were identified, varying from 1 to 15 hybrids per cross, with a median of 3 hybrids per combination. Remarkably, almost all strains could crossbreed with at least one strain from the opposing species, except for the *S. cerevisiae* M9.1 bioethanol strain. The beer clade *S. cerevisiae* strain NCYC_88 demonstrated the highest hybridization success rate among *S. cerevisiae* strains at 52.6% (**Table S3**). In contrast, the *S. cerevisiae* bioethanol M9.1 strain did not produce hybrids with any *S. eubayanus* parental strain. Among the *S. eubayanus* strains, CL248.1 (PB-2 lineage) had the highest hybridization success rate at 38.9% (**Table S3)**, while CL620.1 represented the strain with the highest number of hybrid colonies (31 hybrids, **Figure 1D)**. Notably, the cross between CL620.1 x YJM271 resulted in the highest number of hybrid colonies from a single cross, producing 15 hybrids. These results highlight the substantial viability of generating inter-species hybrids across most *S. cerevisiae* and *S. eubayanus* lineages.

### The hybrid’s fermentative profile is dependent on the S. cerevisiae parental lineage

We evaluated the fermentative capacity of 47 hybrids resulting from 21 different crosses, including their parental strains and the commercial W34/70 strain (**Table S4**). These hybrids exhibited a broad range of CO_2_ production levels, covering the parental phenotypic space with average values spanning from 19.7 to 35 g/L (**Figure 2A, Table S5**). Notably, hybrids sharing the parental *S. cerevisiae* strain NCYC_88 from the beer lineage exhibited the highest CO_2_ production levels (average 31.58 ± 2.17 g/L, *p*-value < 0.05, one-way ANOVA, **Figure S1A**). Building upon this observation, we investigated whether the lineage significantly influenced the hybrids’ fermentative capacity. Indeed, hybrids originating from the *S. cerevisiae* beer lineage consistently showed greater CO_2_ production (31.07 ± 2.21 g/L) compared to hybrids from other lineages (*p*-value < 0.05, Mann-Whitney-Wilcoxon test, **Figure 2B**). Conversely, hybrids from the Wine, Bioethanol, and Sake lineages did not exhibit significant differences in CO_2_ production among themselves (26.64 ± 2.19, 26.73 ± 3.11, and 28.19 ± 0.65 g/L, respectively; *p*-value > 0.05, Mann-Whitney-Wilcoxon test, **Figure 2B**). Interestingly, hybrids stemming from distinct *S. eubayanus* Patagonian clades demonstrated comparable levels of CO_2_ production (*p*-value > 0.05, Mann-Whitney-Wilcoxon test, **Figure 2C, Figure S1B**). Hybrids from the *S. cerevisiae* Beer lineage also exhibited superior maltotriose consumption (56.5 ± 30.38 %, *p*-value < 0.05, ANOVA) compared to hybrids from other lineages (7.0 ± 1.90 %; 6.62 ± 1.19 %; 5.31 ± 1.02 for Sake, Wine and Bioethanol, respectively, **Figure 2D**, **Table S6, Table S7**). Consequently, a positive correlation was observed between maltotriose consumption and the fermentative capacity of these hybrids (R = 0.76, *p*-value < 0.001, Pearson correlation test, **Figure S2**). These results suggest that the fermentation potential of hybrids is predominantly influenced by the *S. cerevisiae* lineage, rather than the *S. eubayanus* parent.

**Figure 2.**
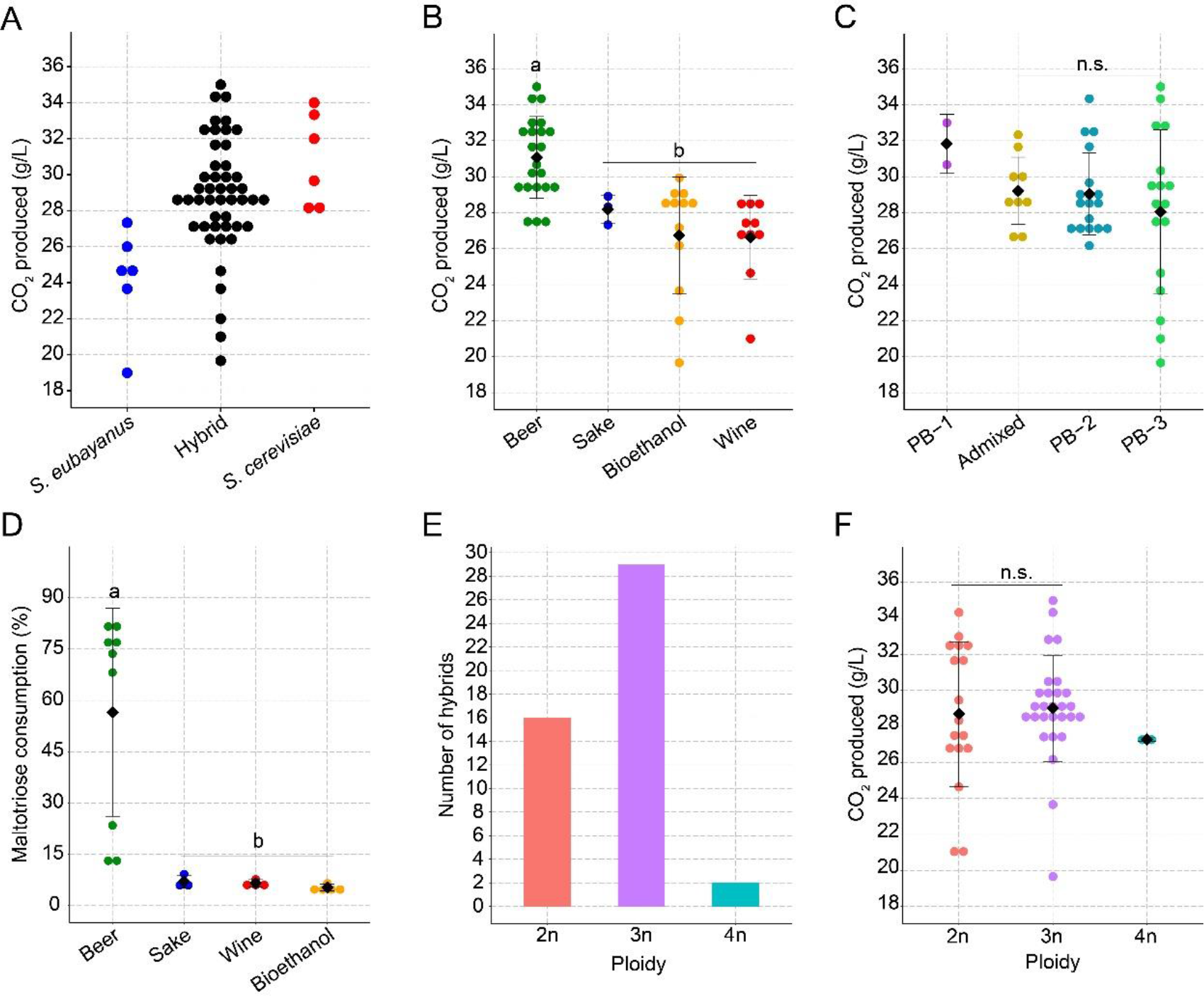
Fermentative capacity of *S. cerevisiae* x *S. eubayanus* hybrids. (A) CO_2_ production levels in *S. eubayanus* (blue), Hybrids (black), and *S. cerevisiae* (red) strains under 12 °P wort. CO_2_ production levels in *de novo* lager hybrids depending on the (B) De novo hybrids grouped by *S. cerevisiae* parental lineage: Beer (green), Sake (blue), Bioethanol (yellow) and Wine (red) lineages and (C) De novo hybrids grouped by *S. eubayanus* lineages: PB-1 (purple), PB-2 (light blue), PB-3 (light green) and admixed (yellow). (D) Maltotriose consumption levels (%) in *de novo* lager hybrids depending on the *S. cerevisiae* parental lineage: Beer (green), Sake (blue), Bioethanol (yellow) and Wine (red) lineages. (E) Ploidy levels determined by FACS in *de novo* lager hybrids after rare mating. (F) CO_2_ production levels depending on the ploidy level. Mean values are depicted by diamonds. Different letters reflect statistically significant differences between strains with a *p*-value < 0.05, one-way ANOVA. n.s. denotes non-significant differences.

Given that the generation of *S. eubayanus* x *S. cerevisiae* hybrids involved rare mating, the resulting hybrids may present diverse ploidy levels potentially associated with brewing-relevant phenotypic traits, such as fermentative capacity. To explore this possibility, we analysed the ploidy levels of the 47 hybrid strains. The observed ploidy spanned from diploid (2n) to tetraploid (4n) (**Figure 2E, Figure S3, Table S8**). Specifically, 16 strains were identified as diploid, 29 as triploid, and 2 displayed a tetraploid state. These results suggest that the rare mating strategy usually involves a 2n x 1n cross. Next, we explored the relationship between ploidy and the fermentative capacity of the 47 hybrids. Upon analyzing these parameters, we did not detect significant differences in the impact of ploidy on CO_2_ production (*p*-value > 0.05, Mann-Whitney-Wilcoxon test, **Figure 2F**). Similarly, when examining the influence of *S. cerevisiae* and *S. eubayanus* lineages and individual parental strains per ploidy level on fermentative capacity, most had no significant differences (*p*-value > 0.05, Mann-Whitney-Wilcoxon test, **Figures S4**). However, the *S. eubayanus* CL248.1 strain exhibited a notable exception, showing higher CO_2_ production in 3n hybrids than 2n hybrids (*p*- value < 0.05, Mann-Whitney-Wilcoxon test, **Figure S4D**). These results suggest that the *S. cerevisiae* lineage is the primary determinant of the hybrid’s fermentative capacity and that ploidy levels might not influence this trait in the lager’s hybrid background.

### S. cerevisiae lineages influence the ethanol and osmotic stress tolerance

To evaluate fitness differences among various *S. cerevisiae* x *S. eubayanus* hybrids, we chose a single representative hybrid per cross combination (21 hybrids). We subjected them to phenotypic assessments and estimated the Area Under the Curve (AUC) under different microculture conditions, including diverse growth temperatures (4°C, 12°C, 20°C, 25°C, 30°C, and 37°C), ethanol tolerance levels (6%, 9%, and 12% v/v), and carbon sources pertinent to the beer brewing process (2% and 20% maltose, 2% maltotriose, and 20 °P wort). Generally, hybrids originating from the Beer lineage exhibited the highest fitness across the tested conditions compared to other lineages, except under 12% ethanol concentration (**Figure 3A**, yellow bar, *p*-value < 0.05, ANOVA, **Table S9**). Conversely, hybrids derived mostly from the Wine lineage demonstrated greater AUC levels under ethanol 12% (**Figure 3B**, green bar, *p*-value < 0.05, ANOVA), representing the only scenario where hybrids from the Beer lineage did not display the highest fitness. Overall, hybrids from the Beer lineage consistently outperformed others, while *S. eubayanus* parental strains exhibited the lowest growth fitness values across conditions, indicating that, in most instances, hybrids inherited the phenotypic profile of *S. cerevisiae* (**Figure 3C)**.

**Figure 3.**
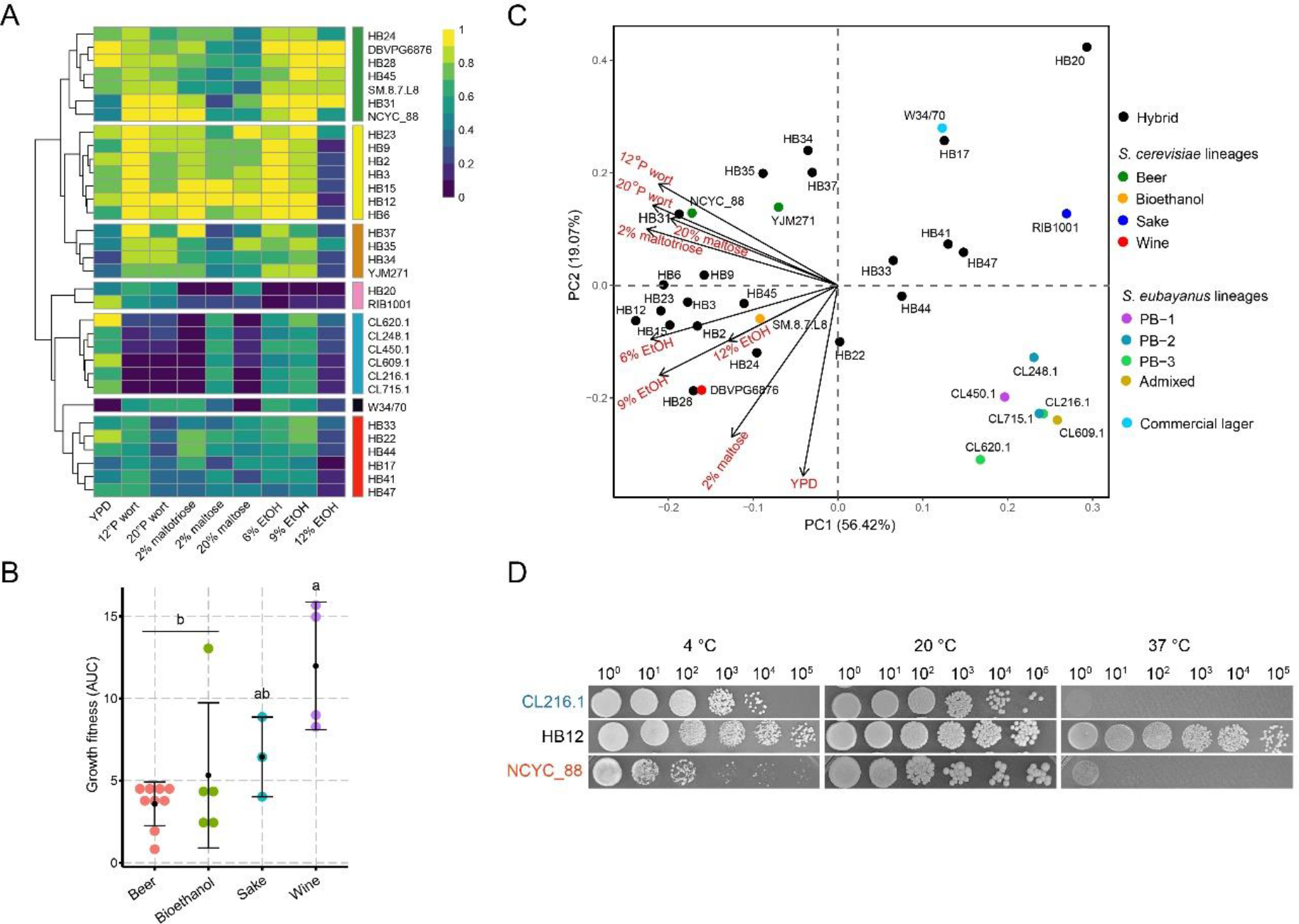
Phenotypic variability across de novo lager hybrids. (A). Heat map depicting the phenotypic diversity in *de novo* lager hybrids obtained from ten assessed conditions. Strains are grouped by hierarchical clustering from AUC data, and names & colours highlight groups of hybrids with similar phenotypes. The heat maps were elaborated based on a 0-1 normalization within each phenotype, with 0 and 1 representing the lowest and highest growth values, respectively. (B) AUC levels for 12% ethanol tolerance in different hybrids depending on the *S. cerevisiae* lineage. Black dots depict mean values. (C) PCA depicting the overall association of hybrids and *S. cerevisiae* or *S. eubayanus* parental lineages (D) Plate spotting assay using 10-fold serial dilution of the HB12 hybrids and its corresponding parental strains grown at different temperatures (4°C – 20°C and 37°C).

Subsequently, we investigated yeast growth under various temperature conditions. Hybrids displayed a wider growth range across temperatures than their parental species (**Figure 3D**). Notably, at low temperatures (4°C, **Figure S5**), hybrids exhibited similar or higher fitness than their *S. eubayanus* and *S. cerevisiae* parents, respectively. This pattern persisted at 30°C and 37°C, where hybrids displayed comparable or superior fitness relative to their *S. cerevisiae* and *S. eubayanus* parents, respectively (**Figure S5**).

### The S. cerevisiae parental lineage predominantly determines the hybrid’s volatile compound profile

To explore the novel lager hybrids’ aroma profiles, the 21 hybrids previously assessed were selected to assess the production of volatile compounds (VCs) after beer fermentation. These hybrids showed distinct VC profiles compared to their parental strains, clustering separately (**Figure 4, Table S10**). We identified four distinct hybrid clusters based on their VC profile. The first cluster contained uniquely hybrid HB20 (CL216.1 x YJM271), aromatically distinctive from other hybrids, and characterized by displaying the low levels of most aromas among the strains, except for isobutanol (59.1 ± 1.22 mg/L, sweet, solvent), 2-phenyl ethanol (117.0 ± 3.78 mg/L, roses) and ethyl propanoate (0.77 ± 0.07 mg/L, fruity) (**Figure 4**, yellow bar). The second hybrid’s cluster predominantly steamed from the *S. cerevisiae* NCYC_88 strain (beer clade). However, all these hybrids exhibited a VCs profile similar to their corresponding *S. eubayanus* parental strains, likely reflecting a recessive NCYC_88 strain VC profile inheritance. These hybrids produced low levels of isobutanol (31.6 ± 2.49 to 44.4 mg/L ± 5.34, sweet, solvent), and higher levels of 2-phenylethyl acetate (1.8 ± 0.16 to 2.8 ± 0.15, roses), ethyl acetate (23.77 ± 1.79 to 33.59 ± 2.36, fruity, sweet) and other alcohols, such as 2-phenyl ethanol (100.9 ± 1.94 to 120.15 ± 3.44, roses) and isoamyl alcohol (226.9 ± 17.56 to 280.5 ± 1.97, whiskey, solvent) (**Figure 4**, green bar). The third hybrid’s cluster, which involved strains derived from crosses including the Bioethanol SM.8.7.L8 and Sake RIB1001 *S. cerevisiae* parents, was characterized by the presence of higher levels of ethyl decanoate (0.11 ± 0.01 to 0.238 ± 0.05 mg/L, fruity) and medium-chain fatty acids, including octanoic (4.07 ± 0.58 to 7.76 ± 0.46 mg/L) acid and decanoic acid (1.5 ± 0.64 to 3.4 ± 0.39 mg/L) (waxy or fatty flavors) (**Figure 4**, fuchsia bar). The fourth cluster, primarily emerging from crosses involving the wine DBVPG6876 and beer YJM271 *S. cerevisiae* strains, exhibited lower production levels of most VCs (**Figure 4**, cyan bar).

**Figure 4.**
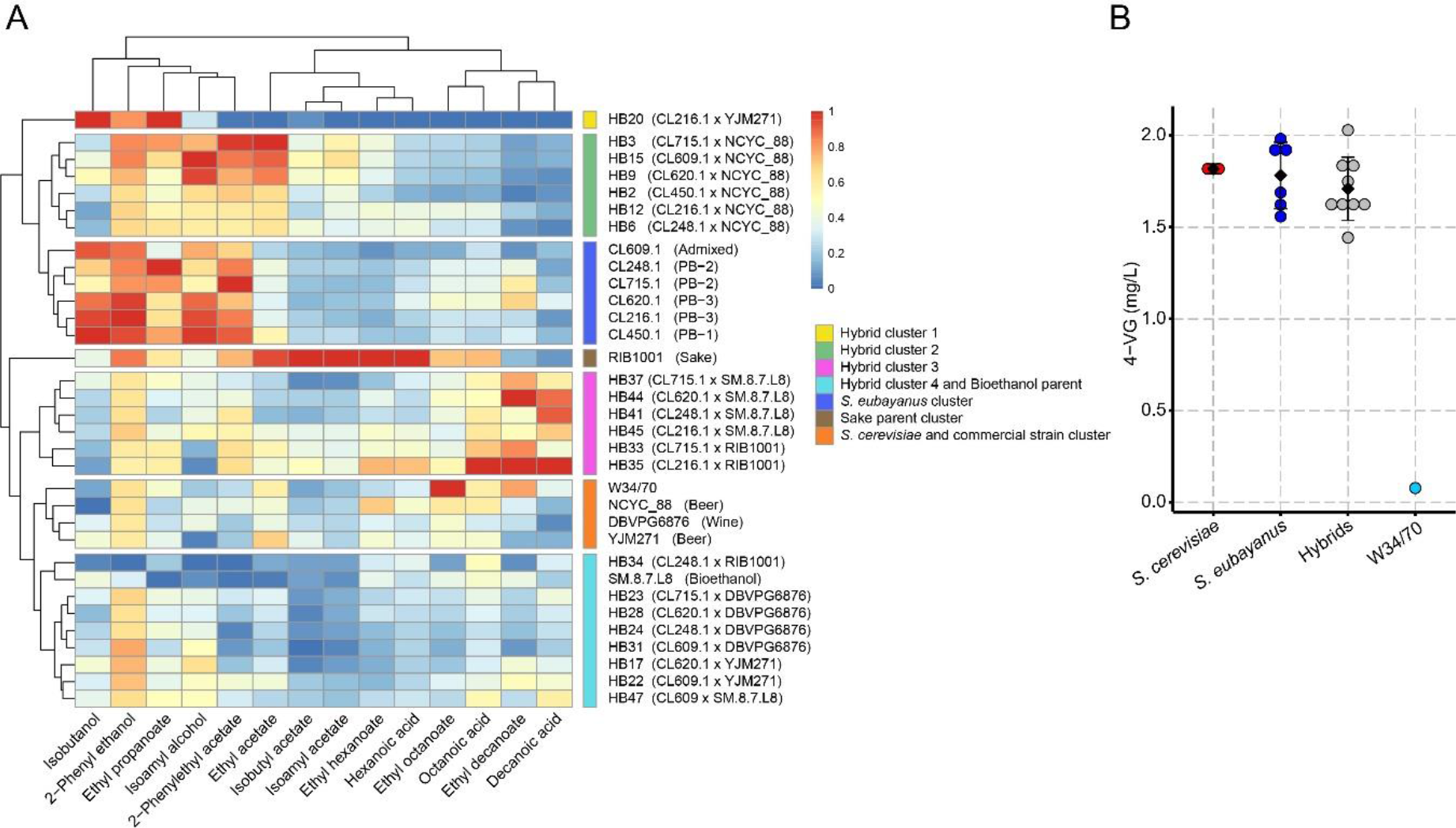
Volatile compound production profile of *de novo* lager hybrids. Beer wort fermentations were carried out for 14 days at 12 °C, and volatile compounds were analyzed by GC-FID at the fermentation endpoint. A) We used a 0-1 normalization within each phenotype, with 0 and 1 representing the lowest and highest VCs values, respectively. Clusters containing strains with similar aroma profiles are highlighted with colored heatmap sidebars. Hybrids’ aroma profiles were classified in four distinctive clusters. B) 4-VG production of hybrids (grey) from Beer and Bioethanol ancestry are represented along with their *S. eubayanus* (blue) and *S. cerevisiae* (red) parental strains. Diamonds depict mean values. No significant differences were detected among hybrids and their parental strains (*p*-value > 0.05, ANOVA). Commercial strain W34/70 (light blue) did not show 4-VG production.

Subsequently, we determined the correlation between VC production in the hybrid backgrounds and the corresponding parental lineages in each species. No significant differences were observed across *S. eubayanus* lineages (*p*-value > 0.05, Mann-Whitney-Wilcoxon test, **Figure S6, Table S11**).

Conversely, hybrids from different *S. cerevisiae* lineages exhibited significant differences in most evaluated aromas (**Figure S7, Table S11**), particularly the brewing lineage, which displayed a fruity and sweet profile with a significantly higher production of ethyl acetate (23,01± 8.39), isoamyl alcohol (236.5 ± 28.0) and 2-phenyl ethanol (109.9 ±7.80) (*p*-value < 0.05, Mann-Whitney-Wilcoxon test, **Figure S7, Table S11**). Hybrids emerging from the beer clade exhibited significantly lower production levels of undesired VC, such as octanoic and decanoic acids (2.1 ± 0.93 and 0.65 ± 0.48, respectively) than hybrids from the bioethanol lineage (4.5 ± 0.35, 2.5 ± 0.51, *p*-value > 0.05, Mann-Whitney-Wilcoxon test, **Figure S7**). Since *S. eubayanus* and some domesticated yeasts produced the phenolic aroma 4-vinyl guaiacol (4-VG), we selected 9 hybrids strains from the Beer and Bioethanol lineages to evaluate its production. We detected similar production levels of 4-VG in hybrids and their corresponding parental strains with no significant differences (*p*-value > 0.05, ANOVA, **Figure 4B, Table S12**), demonstrating a dominant inheritance of this trait.

To evaluate whether novel hybrids displayed enhanced phenotypic traits compared to their parental strains, we assessed best parent heterosis (BPH) across each VC. Our analysis revealed 20 hybrids exhibiting BPH for at least one VC, except for the HB28 hybrid. Interestingly, hybrids derived from the *S. cerevisiae* NCYC_88 strain exhibited 6 out of 14 VCs with BPH, demonstrating the high levels of heterosis in the novel hybrids. BPH hierarchical clustering revealed four distinct hybrid clusters, mostly determined by the *S. cerevisiae* lineage (**Figure 5)**. For example, the third cluster, predominantly included hybrids from the beer parent NCYC_88 and displayed higher BPH values for ethyl acetate (0.14 ± 0.15 to 0.68 ± 0.09, fruity, sweet), isobutyl acetate (0.08 ± 0.04 to 0.75 ± 0.04, fruity, banana), and isoamyl acetate (0.39 ± 0.37 to 1.46 ± 0.19, banana, **Figure 5**, **Table S13**). When examining BPH correlation depending on the *S. eubayanus* parental lineage, significant differences were solely found for ethyl propanoate between PB-2 (-0.32 ± 0.18) and the PB-3 (-0.12 ± 0.17) and Admixed lineages (0.09 ± 0.20) (*p-value* < 0.05, Mann-Whitney-Wilcoxon test, **Figure S8**, **Table S14**). However, in *S. cerevisiae,* we found a significant trend depending on the lineage. In this case, the brewing lineage exhibited significantly greater BPH levels in isoamyl acetate (0.38 ± 0.77) and ethyl propanoate (-0.03 ± 0.21) than Sake (-0.67 ± 0.24) and Wine/European lineages (- 0.27 ± 0.24), respectively (*p*-value < 0.05, Mann-Whitney-Wilcoxon test**, Figure S9, Table S14**). Similarly, hybrids from the bioethanol lineage displayed high BPH values in ethyl hexanoate (-0.13 ± 0.22), ethyl octanoate (-0.07 ± 0.15), ethyl decanoate (0.18 ± 0.29), hexanoic acid (0.07 ± 0.25), octanoic acid (0.20 ± 0.09), and decanoic acid (1.75 ± 0.55) production, mostly distinguishing themselves from brewing and wine lineage strains (*p*-value < 0.05, Mann-Whitney-Wilcoxon test**, Figure S9, Table S13**). These results demonstrate that differences in the hybrid’s VCs profile is predominantly exerted by the *S. cerevisiae* parental lineage rather than being significantly affected by the *S. eubayanus* lineage.

**Figure 5.**
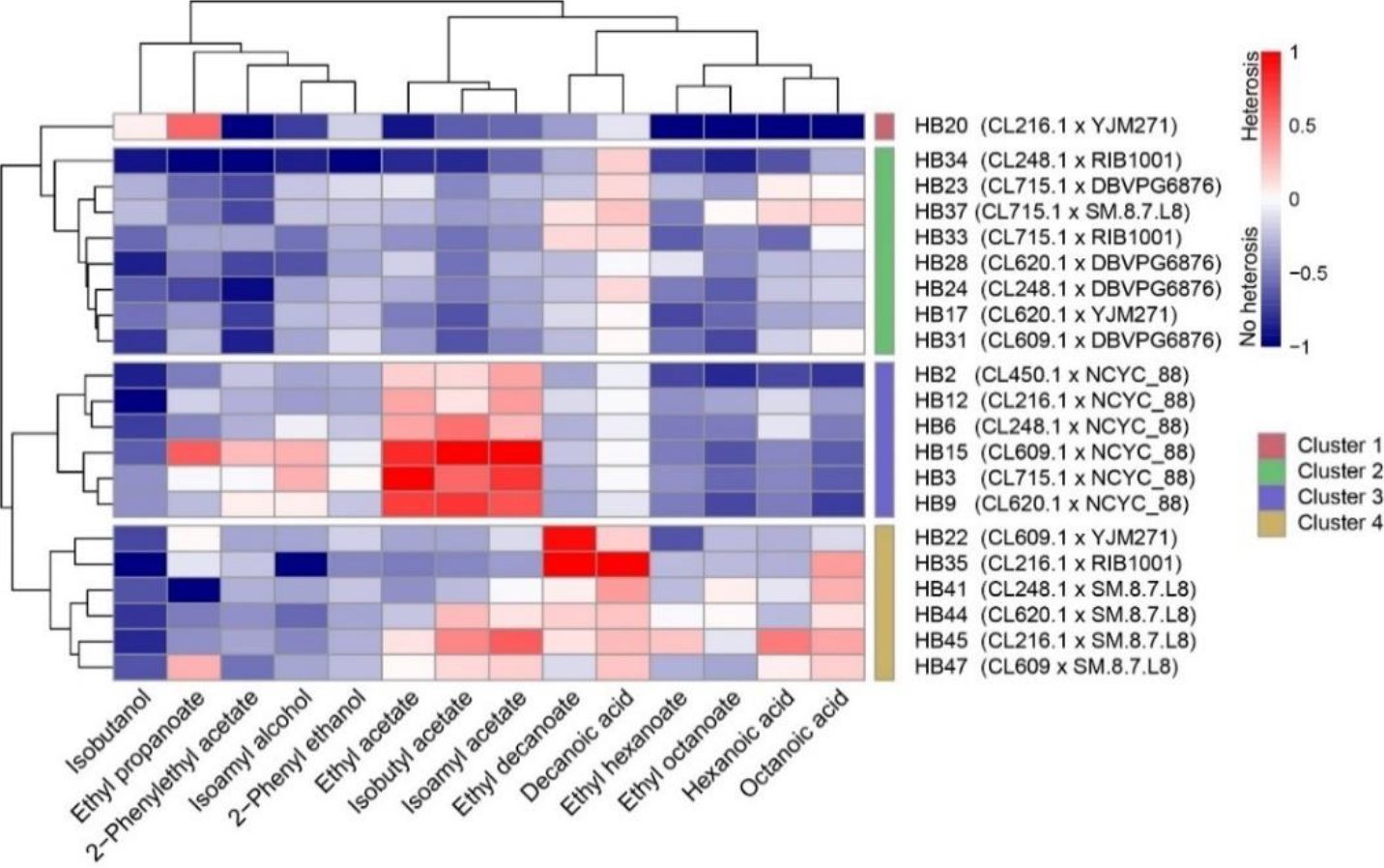
Best Parent Heterosis profile based on volatile compound production in *de novo* lager hybrids. Normalised positive and negative BPH values are depicted on a scale from -1 to 1 relative to the highest absolute BPH value. Hybrids’ BPH profiles were classified into four distinctive clusters.

Finally, exploring the correlation between ploidy and BPH did not detect an overall significant correlation (*p*-value > 0.05, Mann-Whitney-Wilcoxon test**, Figure S10** and **S11**). However, some triploid strains exhibited the highest BPH values and greater variance (CV = 0.580) compared to diploids (CV = 0.295) and tetraploids (CV = 0.186). These results suggest an increased BPH phenotypic variability among triploid hybrids compared to other ploidy levels.

### Gene expression differences between polyploid hybrids

To understand how gene expression underlies the inheritance of brewing traits, we conducted an RNA-seq analysis after 24 hours under beer wort on two *de novo* hybrids characterized by distinct VC profiles exhibiting BPH: HB6 (CL248.1 X NCYC_88) and HB41 (CL248.1 x SM.8.7.L8). By examining mean read counts per gene across chromosomes, we observed a differential representation of parental genomes in each strain, suggesting that HB6 has a greater representation of *S. cerevisiae* (2n *S. cerevisiae* x 1n *S. eubayanus*), while HB41 exhibited an opposite pattern (1n *S. cerevisiae* x 2n *S. eubayanus*) (**Figure S12**). Subsequently, we compared differential gene expression between hybrids. We found 138 genes upregulated in HB6 and 178 in HB41 (log2 fold change > 1, **Figure 6A**, **Table S15**). The metabolic pathways enriched in each hybrid strain, according to KEGG pathways, indicating HB6’s emphasis on 2-oxocarboxylic acid metabolism, while HB41’s exhibited over-expression on pathways such as lysine biosynthesis and the pentose phosphate pathway, which are integral to cellular building blocks and energy metabolism (**Figure 6B**). Similarly, GO terms such as amino acid biosynthetic, organic acid metabolic, and carboxylic biosynthetic processes correlate with the KEGG pathways related to amino acid metabolism highlighted in HB41. In parallel, we observed disparities in carbohydrate metabolism between both hybrids according to GO terms. Specifically, HB6 demonstrated upregulation of genes associated with maltose metabolism, including the maltase *MAL12*, the permease *MAL31*, the transcriptional activator *MAL33*, and the low glucose-induced transporter *HXT14*. In contrast, HB41 exhibited upregulation of genes primarily linked to glucose metabolism, such as *HXT2*, *HXT4*, and *HXT6*. Additionally, HB6 displayed higher gene expression levels in stress tolerance genes, such as *TPS2*, *TSL1*, and *HSP30*. *TPS2* and *TSL1* are implicated in trehalose biosynthesis, while *HSP30* contributes to tolerance against ethanol.

**Figure 6.**
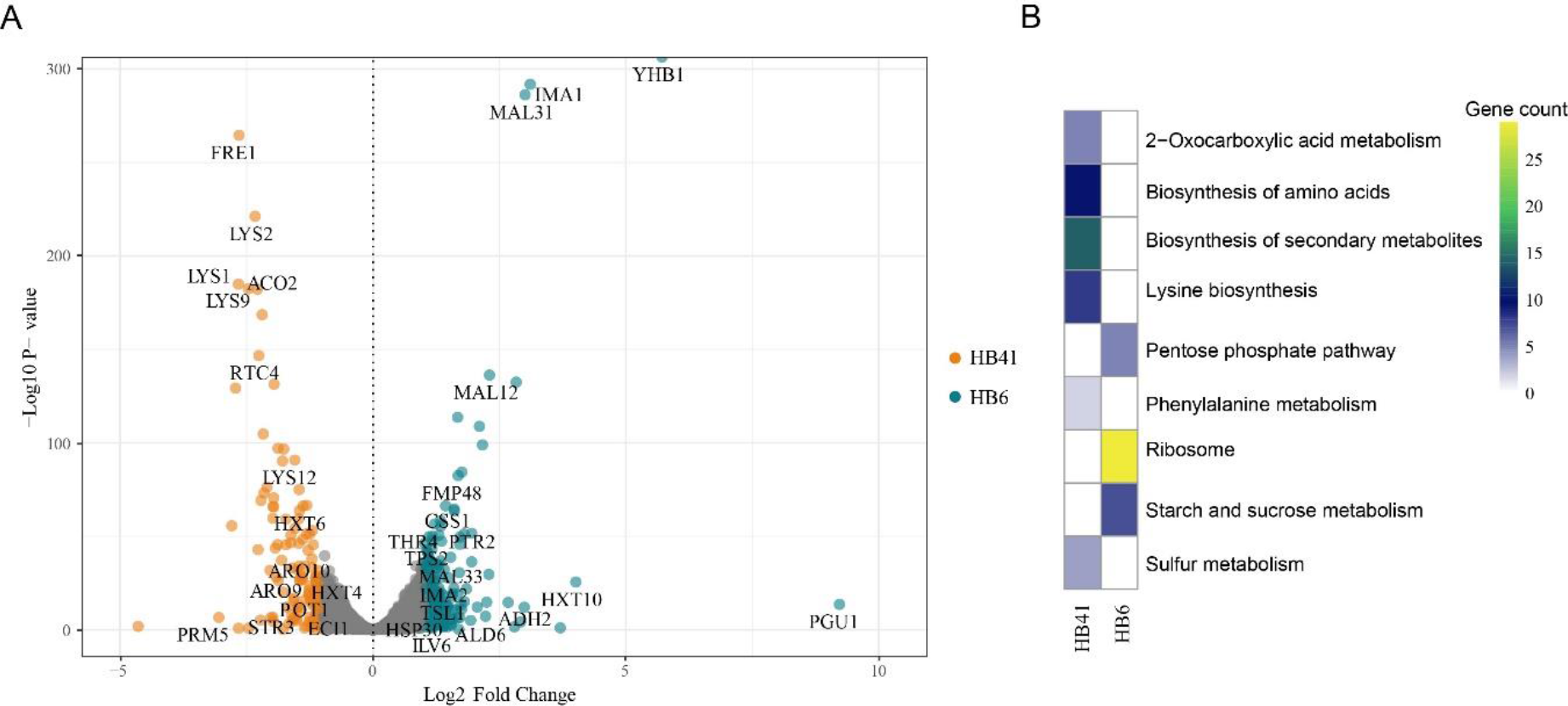
Transcriptome analysis between HB6 and HB41 hybrid strains. (A) Volcano plot depiciting differentially expressed genes (DEGs), and upregulted genes in HB6 (green) and HB41 (orange), hybrid strains. (B) Enriched KEEG pathways in HB6 and HB41 hybrids. Colours depict the number of genes on each category.

Next, we determined the impact of gene expression differences on VC production. For this, we focused on genes and pathways related to HB6 displaying BPH for acetate esters, as representative of other hybrids from the Beer lineage. This analysis highlighted *ADH2* and *ALD6* which showed increased expression levels in HB6. *ADH2* catalyzes the conversion of ethanol to acetaldehyde, a substrate utilized by *ALD6* to produce acetate. Furthermore, we observed upregulation of *ILV6*, a subunit of the acetolactate synthase complex involved in branched-chain amino acid biosynthesis within the mitochondria, serving as precursors for isoamyl acetate and isobutyl acetate. In contrast, HB41, representative of hybrids displaying heterosis in medium-chain fatty acids and their respective ethyl esters showed upregulation in genes associated with fatty acid metabolism such as *POT1*, ECI1, *MGA2*, *IZH4*. *POT1* and *ECI1* are peroxisome genes of β-oxidation. *MAG2* is involved in the synthesis of unsaturated fatty acids and induces the activity of *IZH4*, a membrane protein with a potential role in sterol metabolism. Likewise, in HB41, we observed upregulation of *ARO9* and *ARO10*, genes that participate in the metabolism of aromatic aminoacids and their corresponding acetate esters. Although we didn’t assess the production of aromatic thiols, we detected a higher expression of *STR3* in this strain, a peroxisomal β-lyase implicated in the production of 3-mercaptohexanol (3MH, grapefruit).

## DISCUSSION

Hybridization has played a pivotal role driving evolution across many lineages with immediate phenotypic consequences through the expression of hybrid vigour (Goulet et al., 2017). One example is *S. pastorianus*, the yeast instrumental in lager fermentation, which arose from the hybridization between *S. cerevisiae* and *S. eubayanus* (Gibson et al., 2013; Hutzler et al., 2023). However, our understanding of the *S. cerevisiae* and *S. eubayanus* hybridization success remains limited, largely due to the lack of a wide range of *S. eubayanus* parental strains. Previous studies have only utilized a single *S. eubayanus* parental strain, constraining our understanding of the hybridization process (Hebly et al., 2015; Krogerus et al., 2015, 2016; Nikulin et al., 2018). The recent identification of diverse *S. eubayanus* lineages in the Patagonian region presents an opportunity to partly elucidate the molecular underpinnings driving the robust hybrid vigour observed in *S. pastorianus* (Langdon et al., 2020; Nespolo, et al., 2020).

Polyploid hybrids can be generated through the rare mating technique (Morard et al., 2020; Pérez et al., 2022), representing the most successful strategy to generate artificial hybrids in the *Saccharomyces* genus (Lairón-Peris et al., 2020; Pérez et al., 2022; Pérez-Través et al., 2012). This process exploits yeast strains that could have undergone loss of heterozygosity in the genes responsible for sexual pheromones (*MAT*α and *MAT*a), which are situated on chromosome III in both species. Such a loss enables a diploid *MAT*α/*MAT*a strain to transition into either a *MAT*α or *MAT*a diploid state, allowing them to mate with yeasts of the opposite mating type (Lairón-Peris et al., 2020; Morard et al., 2020). To quantify the degree of hybridization success between *S. cerevisiae* and *S. eubayanus* and its consequential impact on fermentation performance and the profile of volatile compounds in newly formed lager hybrids, we meticulously selected parental for rare mating from candidates spanning a diverse array of lineages from both species (Nespolo, et al., 2020; Peter et al., 2018). Our results indicate that hybridization success between diploid *S. eubayanus* and *S. cerevisiae* is common and particularly high in individual lineages, depending on the species. We identified parental strains from the Beer and Bioethanol *S. cerevisiae* lineages and the PB-2 *S. eubayanus* lineage, exhibiting the highest hybridization rates. However, it is noteworthy that no hybrids were produced from one Bioethanol lineage strain. While we did not investigate the underlying reasons for this observation, we believe that certain genetic incompatibilities identified in interspecific hybridizations, particularly those related to mitochondrial and nuclear interactions, might account for the disparities in the mating rates of our hybrids (Blanckaert & Payseur, 2021; Lee et al., 2008; Moran et al., 2024; Swamy et al., 2022). Ploidy analysis revealed that rate mating led to a higher proportion of triploids hybrids. The observed pattern likely arises from two distinct mating scenarios: the mating between spores and diploid parents, which gives rise to triploids, and the mating between haploid spores, resulting in diploids (Krogerus et al., 2015; Nikulin et al., 2018; Pérez et al., 2022). Several studies have linked yeast ploidy levels to their fitness under varying conditions. For instance, Krogerus et al. (2016) reported that a tetraploid *de novo* lager hybrid displayed a higher fermentative capacity than a triploid and diploid counterpart. However, this study only limited set of hybrids. In contrast, Peter et al. (2018) found that *S. cerevisiae* diploids exhibited superior fitness compared to triploids and tetraploids strains. Interestingly, we did not observe a correlation between ploidy and fermentative capacity across our interspecific hybrids. This suggests that ploidy might not be the main factor driving fermentative capacity vigour in *de novo* lager hybrids.

Hybridization can generate hybrids with novel traits compared to parental species, which often can meet the requirements of the fermented food industry, such as high fermentative capacity in wort (Krogerus et al., 2015). The set of hybrids covered the complete phenotypic space comprised of parental species. However, no significant heterosis levels were observed. In this sense, the performance of hybrids emerging from the Beer lineage inherited the maltotriose consumption capacity from the corresponding *S. cerevisiae* parent. Maltotriose is a trisaccharide representing the second most abundant sugar source in the wort, serving as a crucial substrate for yeast metabolism and overall fermentation efficiency (Zastrow et al., 2000). Indeed, we observed *MAL* genes to be upregulated in the HB6 strain, which contained a 2n and 1n composition of *S. cerevisiae* and *S. eubayanus* subgenomes, respectively, compared to the HB41 hybrid. In contrast, HB41, carrying an opposite subgenome pattern, exhibited upregulation of genes associated with glucose metabolism. Our results agree with previous studies demonstrating that interspecific hybrids, including the *S. cerevisiae* Beer lineage, perform better in fermentation than wine hybrids (Mertens et al., 2015). Beer ale strains have been historically selected as the preferred yeast to generate hybrid offspring with increased maltotriose consumption (Mukai et al., 2001; Nikulin et al., 2018). Certain hybrid combinations of negative maltotriose strains could increase the metabolization of this trisaccharide, revealing subgenomes crosstalk (Brouwers et al., 2019). However, our study did not evidence hybrid vigour, possibly attributable to inefficient maltotriose transporters in the parental strains or a lack of crosstalk between the subgenomes for this trait.

The current ambition to develop innovative lager yeast strains seeks to modernise aroma profiles in lager beers. In this context, hybridization emerges as a promising strategy for creating yeasts with distinctive aroma profiles (Giannakou et al., 2021; Krogerus et al., 2016; Mertens et al., 2015; Pérez et al., 2022) . Previously, the volatile compound profiles in hybrids have been associated with ploidy (Krogerus et al., 2016). We generally did not identify a significant correlation between the hybrid’s ploidy and VCs best parent heterosis. Overall, these results indicate that ploidy might not significantly affect brewing traits. Instead, when evaluating the contribution of the species’ parental lineages, we found that the hybrid’s volatile compound profile primarily differs depending on the *S. cerevisae* parental lineage rather than the *S. eubayanus* lineage. Most notably, hybrids emerging from the Beer and Bioethanol lineages differentiated each other in the production of acetate esters and higher alcohols (fruity/flowery), fatty acids (waxy), and their derivative esters (flowery). Although waxy aromas are considered off-flavours in beer, octanoic and decanoic acids in our Bioethanol hybrids were detected within the range of commercial beverages (Ferreira et al., 2019; Siebert, 1999). The volatile compound profile of *S. cerevisiae* Beer hybrids varied depending on the *S. eubayanus* parent, suggesting that the volatile compound machinery of *S. eubayanus* exhibits a dominant inheritance over that of the Beer *S. cerevisiae* strain in shaping these traits. Inherited phenotypes from one parental strain have been documented in hybrid studies (Catallo et al., 2021; Giannakou et al., 2021; Kanter et al., 2020). In this context, the superior parent heterosis in volatile compound production may stem from the metabolic rewiring in interspecific hybrids (Herbst et al., 2017).

Our VC pattern analysis identified two hybrid groups with distinct heterotic aroma profiles. Hybrids from beer ancestry demonstrated high heterosis values for ethyl acetate, isobutyl acetate, and isoamyl acetate. Within this group, the *de novo* hybrid HB6 exhibited increased gene expression levels of alcohol metabolism genes, such as *ADH2* and *ALD6*, which catalyse acetate synthesis (Simpson-Lavy & Kupiec, 2019). The acetyl-CoA form of acetate is a precursor for the aforementioned acetate esters (Holt et al., 2019). In this sense, the over-expression of *ALD6* has been used to increase the production of ethyl acetate under fermentation conditions (Shi et al., 2021). In addition, we found *ILV6* upregulated in HB6, a subunit of the complex acetolactate synthase that increases the activity of *ILV2* in branched-chain amino acid biosynthesis in mitochondria (Takpho et al., 2018). Precursors of this pathway are used to synthesise isobutyl acetate, and isoamyl acetate (Holt et al., 2019). On the contrary, hybrids from Bioethanol ancestry, such as HB41, exhibited heterosis for octanoic acid, decanoic acid, and their respective ethyl esters. The excretion of these compounds into the media has been associated with detoxification mechanisms (Borrull et al., 2015; Saerens et al., 2010). Previous studies have shown that ethanol enhances the toxicity of octanoic and decanoic acids within the cell (Legras et al., 2010). In this sense, expression analysis in HB41, evidenced the upregulation of genes involved in β-oxidation such as *POT1* and *ECI1*(Rajvanshi et al., 2017). In this way, hybrid strains derived from the *S. cerevisiae* Bioethanol lineage might possess a heightened fatty acid metabolism as a response to tolerate the alcoholic environment. For instance, HB41 exhibited upregulation in the gene expression of *MGA2* and *IZH4*, both associated with ergosterol biosynthesis, a well-known strategy that yeasts employ to tolerate alcoholic stress (Lairón-Peris et al., 2021).

In conclusion, our study emphasizes the importance of genetic diversity in the parental strains background, as demonstrated by the varied hybridization success rates and phenotypic outcomes when different lineages of *S. cerevisiae* and *S. eubayanus* are crossed. Interestingly, we revealed that ploidy levels among the hybrids might not be the primary factor influencing fermentative performance. Instead, we identified a significant relationship between the specific parental lineage and the volatile compound and fermentative profiles of the hybrids, indicating lineage-specific inheritance of traits that are crucial for the brewing industry. Future research should focus on understanding the genetic and molecular basis of hybrid traits to optimize yeast strains for specific brewing requirements, potentially leading to the development of yeasts with tailor-made characteristics for improved fermentation and aroma profiles.

## ACKNOWLEDGMENTS

We acknowledge Fundación Ciencia & Vida for providing infrastructure, laboratory space, and experiment equipment. This research was partially supported by the supercomputing infrastructure of the National Laboratory for High Performance Computing Chile (NLHPC, ECM-02).

## ETHICAL STATEMENT

This article does not contain any studies with human nor animal subjects performed by any of the authors.

## DATA AVAILABILITY

All fastq sequences were deposited in the National Center for Biotechnology Information (NCBI) as a Sequence Read Archive under the BioProject accession number PRJNA1103204.

## SUPPLEMENTARY INFORMATION FIGURE LEGENDS

Figure S1. Fermentative profile of *S. eubayanus* x *S. cerevisae* hybrids depending on the parental strain. CO_2_ production levels in *de novo* lager hybrids in 12 °P wort depending on the (A) *S. cerevisiae* and (B) *S. eubayanus* parental strains. Lineage are shown in different colours. Mean values are depicted by diamonds. Statistical differences were calculated by using Wilcoxon–Mann– Whitney test, p-value < 0.05. n.s. denotes non-significant differences.

Figure S2. **Hybrid’s maltotriose consumption levels under beer wort.** (A) Maltotriose consumption in *de novo* lager hybrids depending on the *S. eubayanus* parental lineages. Mean values are depicted by diamonds. Statistical differences were calculated by using Wilcoxon–Mann– Whitney test, p-value < 0.05. n.s. denotes non-significant differences. (B) Correlation between maltotriose consumption and fermentative capacity in 21 de novo lager hybrids representative from each cross combination (p-value < 0.001, Pearson correlation test).

Figure S3. **Ploidy levels of *de novo* hybrids.** Ploidy levels were determined using flow cytometry. Fluorescence curves depicting the DNA content of each strain labeled with propidium iodide are provided. The fluorescence intensity is represented on the x-axis, while the y-axis denotes cell counts, in the y-axis. Approximately 150,000 single cells were used for determining ploidy. Haploid, diploid, and tetraploid standards (S. cerevisiae YPS128-*Mat* a , S. cerevisiae DBVPG6876, and S. pastorianus W34/70 strains, respectively) were employed for calibration.

Figure S4. **Fermentative profile of *de novo* lager hybrids sorted by parental lineages and ploidy number.** CO_2_ production levels in *de novo* lager hybrids depending on the (A) *S. cerevisiae* lineage, (B) *S. cerevisiae* strain, (C) *S. eubayanus* lineages and (D) *S. eubayanus* strain. Also, the same analysis was determined depending on the S. cerevisiae (C) and S. eubayanus (D) parental strains. Mean values are depicted by diamonds. Statistical differences were calculated by using Wilcoxon–Mann–Whitney test, p-value < 0.05.

Figure S5. **Hybrid’s temperature tolerance.** A serial dilution assay was carried out on YPD plates with incubation temperatures ranging from 4 to 37 °C. *S. eubayanus* and *S. cerevisiae* parental strains are indicated in blue and red, respectively. Additionally, commercial lager strain W34/70 was included in the assay. Photographs of temperature plate sets were taken at different times (see Methodology for details).

Figure S6. **Volatile compounds production of hybrids depending on the *S. eubayanus* parental lineages.** Beer wort fermentations were conducted for 14 days at 12 °C, and volatile compounds at the end of the fermentation process were determined by GC-FID chromatography. Mean values are depicted by diamonds. No statistical differences were observed in the analysis (Wilcoxon–Mann– Whitney test, p-value > 0.05).

Figure S7. **Volatile compounds production of hybrids depending on the *S. cerevisiae* parental lineages.** Beer wort fermentations were conducted for 14 days at 12 °C, and volatile compounds at the end of the fermentation process were estimated by GC-FID chromatography. Mean values are depicted by diamonds. No statistical differences were observed in the analysis (Wilcoxon–Mann– Whitney test, p-value > 0.05).

Figure S8. **Volatile compound heterosis levels in hybrids depending on the *S. eubayanus* parental lineage.** Heterosis, expressed as best parent heterosis (BPH) was determined relative to the volatile compound production levels of the best parent for the corresponding trait. Red dashed line over the 0 value represents the for the corresponding trait level of the best parent of each hybrid. Mean values are depicted by diamonds. Statistical differences were detected only in ethyl propanoate (Wilcoxon–Mann–Whitney test, p-value < 0.05).

Figure S9. **Volatile compound heterosis in hybrids depending on the *S. cerevisiae* lineage**. Heterosis, expressed as best parent heterosis (BPH) was determined relative to the volatile compound production levels of the best parent for the corresponding trait. Red dashed line over the 0 value represents the volatile compound production levels of the best parent for each hybrid. Mean values are depicted by diamonds. Statistical differences were detected in 13 out of 14 aromas (Wilcoxon–Mann–Whitney test, p-value < 0.05).

Figure S10. **Influence of ploidy on volatile compound heterosis**. Diploid (red), triploid (purple) and tetraploid (sky blue) hybrids are shown. All BPH values for fourteen volatile compounds were included in this analysis. Red dashed line over the 0 value represents the aroma production level of the best parent of each hybrid. Mean values are depicted by diamonds. No statistical differences were observed in the analysis (Wilcoxon–Mann–Whitney test, p-value > 0.05).

Figure S11. **Influence of ploidy on each volatile compound heterosis.** BPH values for fourteen volatile compounds in diploid (red) and triploid (purple) strains are shown. Red dashed line over the 0 value represents the volatile compound levels of the best parent of each hybrid. Mean values are depicted by diamonds. Statistical differences were detected only in ethyl decanoate (Wilcoxon– Mann–Whitney test, p-value < 0.05).

Figure S12. **Gene expression levels depending on the parental allele in HB6 and HB41 hybrids.**

## TABLE LEGENDS

Table S1. Strains used in this study.

Table S2. Sugars consumption (g/L) after 14 days under beer wort fermentation. R= replicates, SD= standar deviation. A. *S. eubayanus* strains, B. *S. cerevisiae*.

Table S3. Hybridization success rate.

Table S4. S. cerevisiae x S. eubayanus hybrids.

Table S5. Fermentative capacity of hybrids under beer wort. R= replicates, SD= standar deviation.

Table S6. Sugars consumption of hybrids after 14 days in beer wort fermentation. R= replicates, SD= standar deviation.

Table S7. Maltoriose consumption in hybrids after 14 days in beer wort fermentation. SD= standar deviation.

Table S8. Ploidy levels in hybrids determined by flow cytometry.

Table S9. Area Under the Curve (AUC) in hybrids grown under different environmental conditions.

Table S10. Volatile compounds production in hybrids and parental strains.

Table S11. Volatile compounds production in hybrids depending on the parental lineage. Table S12. 4-VG production in hybrids and parental strains.

Table S13. Best patent heterosis (BPH) levels in hybrids.

Table S14. Best parent heterosis (BPH) in volatile compounds levels depending on the parental lineage.

Table S15. RNA-seq results. Design = HB6 vs HB41.

